# SMRT long reads and Direct Label and Stain optical maps allow the generation of a high-quality genome assembly for the European barn swallow (*Hirundo rustica rustica*)

**DOI:** 10.1101/374512

**Authors:** Giulio Formenti, Matteo Chiara, Lucy Poveda, Kees-Jan Francoijs, Andrea Bonisoli-Alquati, Luca Canova, Luca Gianfranceschi, David Stephen Horner, Nicola Saino

## Abstract

**Background:** The barn swallow (*Hirundo rustica*) is a migratory bird that has been the focus of a large number of ecological, behavioural and genetic studies. To facilitate further population genetics and genomic studies, here we present a reference genome assembly for the European subspecies (*H. r. rustica*).

**Findings:** As part of the Genome10K (G10K) effort on generating high quality vertebrate genomes, we have assembled a highly contiguous genome assembly using Single Molecule Real-Time (SMRT) DNA sequencing and several Bionano optical map technologies. We compared and integrated optical maps derived both from the Nick, Label, Repair and Stain and from the Direct Label and Stain (DLS) technologies. As proposed by Bionano, the DLS more than doubled the scaffold N50 with respect to the nickase. The dual enzyme hybrid scaffold led to a further marginal increase in scaffold N50 and an overall increase of confidence in the scaffolds. After removal of haplotigs, the final assembly is approximately 1.21 Gbp in size, with a scaffold N50 value of over 25.95 Mbp.

**Conclusions:** This high-quality genome assembly represents a valuable resource for further studies of population genetics and genomics in the barn swallow, and for studies concerning the evolution of avian genomes. It also represents one of the very first genomes assembled by combining SMRT long-read sequencing with the new Bionano DLS technology for scaffolding. The quality of this assembly demonstrates the potential of this methodology to substantially increase the contiguity of genome assemblies.

## Data Description

### Context

The barn swallow is a passerine bird with at least eight recognized subspecies in Europe, Asia and North America. The European barn swallow (*Hirundo rustica rustica*) (Figure 1) breeds in a broad latitudinal range, between 63-68°N and 20-30°N [1]. Numerous evolutionary and ecological studies have focused on its biology, including its life history, sexual selection, and response to climate change. More recently, the barn swallow has become the focus of genetic studies on the divergence between subspecies and populations [2–4] and on the control of phenological traits [5–8]. Due to its synanthropic habits and its cultural value, the barn swallow is also a flagship species in conservation biology [1]. The availability of high-quality genomic resources, including a reference genome, is thus pivotal to further boost the study and conservation of this species.

**Figure 1:**
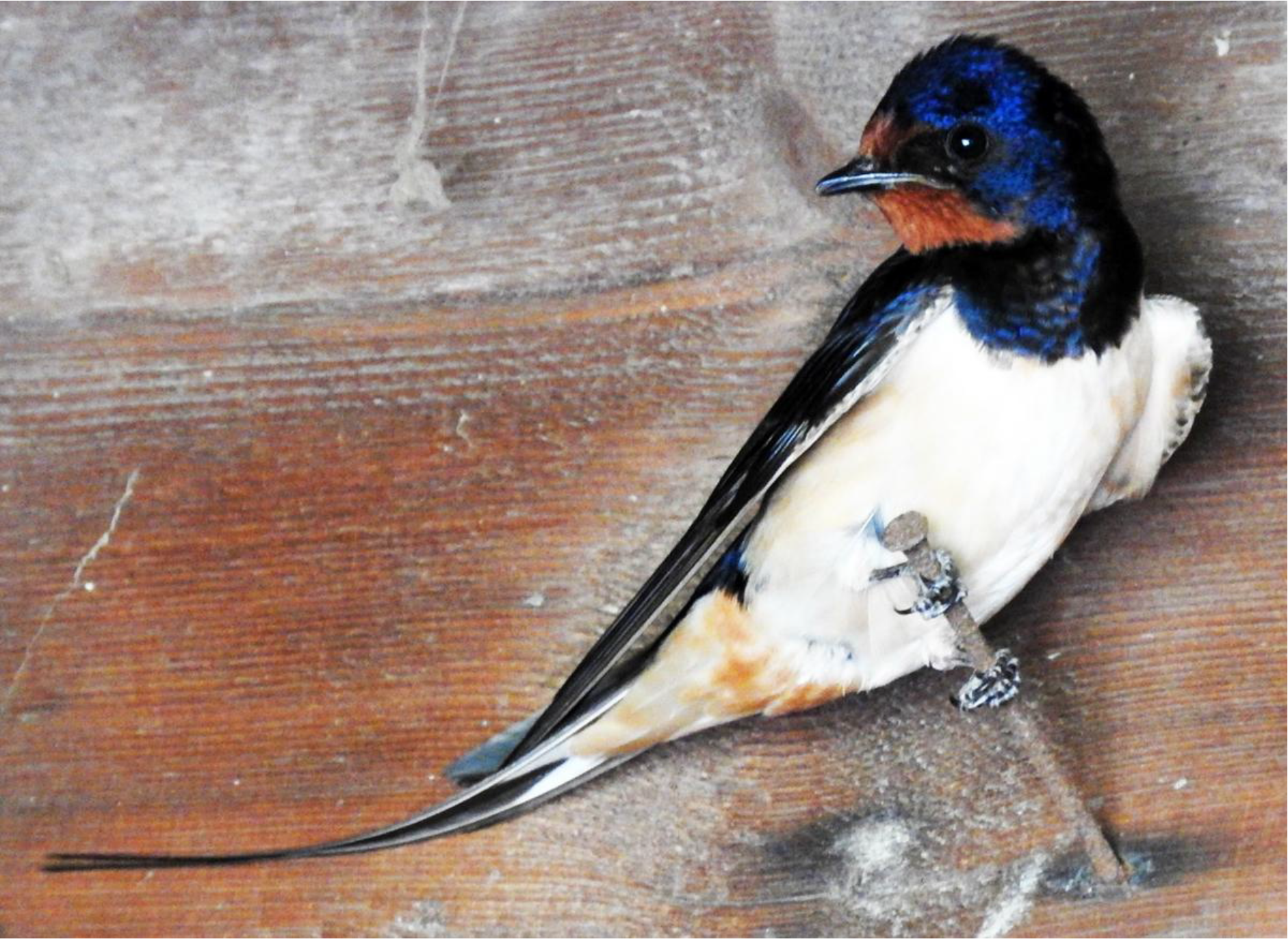
the European barn swallow (*Hirundo rustica rustica*). Courtesy of Chiara Scandolara.

In 2016, Safran and coworkers reported the first draft of the genome for the American subspecies (*Hirundo rustica erythrogaster*) constructed from Illumina paired-end reads [2]. However, it has not been possible to analyze this assembly as neither the raw nor the assembled data were publicly available at the time of preparation of the current manuscript [2].

Here we have employed two single-molecule technologies, Single Molecule Real Time (SMRT) Third-Generation Sequencing (TGS) from Pacific Biosciences (Menlo Park, California, USA) and optical mapping from Bionano Genomics (San Diego, California, USA), to produce a state-of-the-art high-quality genome assembly for the European subspecies. For optical mapping we labelled DNA molecules both with one of the original Nick, Label, Repair and Stain (NLRS) nickases (enzyme Nb.BssSI) and with the new Direct Label and Stain (DLS) approach (enzyme DLE-1). The latter technique was officially released in February 2018 and avoids nicking and subsequent cleavage of DNA molecules during staining [9]. We show that DLS allows a considerable improvement of scaffold contiguity with respect to the nickase tested, consistent with Bionano’s claim. Furthermore, the “dual enzyme” approach affords additional support for scaffold junctions. This genome assembly is among the first to incorporate DLS and SMRT sequencing data, providing assembly contiguity metrics well in excess of those specified for “Platinum genomes” by the Vertebrate Genomes Project (VGP) [10,11]. While this article was under review, the Vertebrate Genomes Project released 15 genome assemblies that incorporate SMRT and DLS data among others, including the hummingbird and Kakapo, with comparable results (Bioproject PRJNA489243) [12].

### Blood sample collection

The blood used as a source of DNA was derived from a minimally invasive sampling performed on a female individual of approximately two years of age during May 2017 in a farm near Milan, in Northern-Italy (45.4N 9.3E). Blood was collected in heparinized capillary tubes. Three hours after collection, the sample was centrifuged to separate blood cells from plasma, and then stored at −80°C.

### DNA extraction and quality control for SMRT library preparation

DNA extraction was performed on blood cells portion of centrifuged whole blood containing nucleated erythrocytes and leukocytes with the Wizard genomic DNA purification kit (Promega, Cat. No. A1125), using the protocol for tissue (not human blood). This kit employs a protocol similar to the classical Phenol/Chloroform DNA extraction, with no vortexing steps after cell lysis. After purification, DNA quality and concentration were assessed by Nanodrop (Thermo Fisher Scientific, Cat. No. ND-1000), and subsequently by Pulsed Field Gel Electrophoresis (PFGE). Detectable DNA was over 23 kbp in size, with the vast majority over 50 kbp and even over 200 kbp (Supplementary Figure 1). PFGE quality results were further confirmed by capillary electrophoresis on FEMTO Pulse instrument (AATI, Cat. No. FP-1002-0275) (Supplementary Figure 2). DNA was stored at −80°C and shipped to the sequencing facility on dry ice.

### SMRT library preparation and sequencing

SMRTbell Express Template Prep Kit (Pacific Biosciences, Cat. No. 101-357-000) was used to produce the insert library. Input genomic DNA (gDNA) concentration was measured on a Qubit Fluorometer dsDNA Broad Range (Life Technologies, Cat. No. 32850). 10 μg of gDNA was mechanically sheared to an average size distribution of 40-50 kbp, using a Megaruptor Device (Diagenode, Cat. No. B06010001). FEMTO Pulse capillary electrophoresis was employed to assess the size of the fragments. 5 μg of sheared gDNA was DNA-damage repaired and end-repaired using polishing enzymes. Blunt-end ligation was used to create the SMRTbell template. A Blue Pippin device (Sage Science, Cat. No. BLU0001) was used to size-select the SMRTbell template and enrich for fragments > 30 kbp, excluding the first two cells for which the library was enriched for fragments > 15 kbp. The size-selected library was checked using FEMTO Pulse and quantified on a Qubit Fluorometer. A ready to sequence SMRT bell-Polymerase Complex was created using the Sequel binding kit 2.0 (Pacific Biosciences, Cat. No. 100-862-200). The Pacific Biosciences Sequel instrument was programmed to sequence the library on 18 Sequel SMRT Cells 1M v2 (Pacific Biosciences, Cat. No. 101-008-000), taking one movie of 10 hours per cell, using the Sequel Sequencing Kit 2.1 (Pacific Biosciences, Cat. No. 101-310-400). After the run, sequencing data quality was checked via the PacBio SMRT Link v5.0.1 software using the “run QC module”. An average of 3.7 Gbp (standard deviation: 1.7) were produced per SMRT cell (average N50 = 25,622 bp), with considerable improvements between the average 15 kbp library and the 30 kbp library (see Supplementary Figure 3 for more detailed statistics). We observed a wide distribution in the GC content of reads (Supplementary Figure 4). This is likely explained by the presence in avian genomes of three classes of chromosomes: macrochromosomes (50-200 Mbp, 5 in chicken), intermediate chromosomes (20-40 Mbp, 5 in chicken) and microchromosomes (12 Mbp on average, 28 in chicken). These last account for only 18% of the total genome but harbour ~31% of all chicken genes, have higher recombination rates and higher GC contents on average [13].

### Assembly of SMRT reads

The final assembly of long reads was conducted with software CANU v1.7 [14] using default parameters except for the “correctedErrorRate” which was set at 0.075. The assembly processes occupied 3,840 CPU hours and 2.2 Tb of RAM for read correction, 768 CPU hours and 1.1 Tb of RAM for the trimming steps, and 3280 CPU hours and 2.2 Tb of RAM for the assembly phase. The long-read assembly contained 3,872 contigs with a N50 of 5.2 Mbp for a total length of 1311.7 Mbp (Table 1 and Supplementary Table 1). Final polishing was performed using the Arrow v2.10 software (Pacific Biosciences) and resulted in final coverage of 45.4x.

### Cell count and DNA extraction for optical mapping

High-molecular weight (HMW) DNA was extracted from 7-8 μl of the cell portion from the same blood sample used for SMRT sequencing with the Blood and Cell Culture DNA Isolation kit (Bionano Genomics, Cat. No. RE-016-10). HMW DNA was extracted by embedding cells in low melting temperature agarose plugs that were incubated with Proteinase K (Qiagen, Cat. No. 158920) and RNAseA (Qiagen, Cat. No. 158924). The plugs were washed and solubilized using Agarase Enzyme (Thermo Fisher Scientific, Cat. No. EO0461) to release HMW DNA and further purified by drop dialysis. DNA was homogenised overnight prior to quantification using a Qubit Fluorometer.

### *In silico* digestion

The genome assembly obtained with CANU was *in silico* digested using Bionano Access software to test whether the nicking enzyme (Nb.BssSI), with recognition sequence (CACGAG), and the non-nicking enzyme DLE-1, with recognition sequence (CTTAAG), were suitable for optical mapping in our bird genome. An average of 16.9 nicks/100 kbp with a nick-to-nick distance N50 of 11,708 bp were expected for Nb.BssSI, while DLE-1 was found to induce 19.1 nicks/100 kbp with a nick-to-nick distance N50 of 8,775 bp, both in line with manufacturer’s requirements.

### DNA labeling for optical mapping

For NLRS, DNA was labelled using the Prep DNA Labeling Kit-NLRS according to manufacturer’s instructions (Bionano Genomics, Cat. No. 80001). 300 ng of purified gDNA was nicked with Nb.BssSI (New England Biolabs, Cat. No. R0681S) in NEB Buffer 3. The nicked DNA was labelled with a fluorescent-dUTP nucleotide analog using Taq DNA polymerase (New England BioLabs, Cat. No. M0267S). After labeling, nicks were ligated with Taq DNA ligase (New England BioLabs, Cat. No. M0208S) in the presence of dNTPs. The backbone of fluorescently labelled DNA was counterstained overnight with YOYO-1 (Bionano Genomics, Cat. No. 80001).

For DLS, DNA was labelled using the Bionano Prep DNA Labeling Kit-DLS (Cat. No. 80005) according to manufacturer’s instructions. 750 ng of purified gDNA was labelled with DLE labeling Mix and subsequently incubated with Proteinase K (Qiagen, Cat. No. 158920) followed by drop dialysis. After the clean-up step, the DNA was pre-stained, homogenised and quantified using on a Qubit Fluorometer to establish the appropriate amount of backbone stain. The reaction was incubated at room temperature for at least 2 hours.

### Generation of optical maps

NLRS and DLS labelled DNA were loaded into a nanochannel array of a Saphyr Chip (Bionano Genomics, Cat. No. FC-030-01) and run by electrophoresis each into a compartment. Linearized DNA molecules were imaged using the Saphyr system and associated software (Bionano Genomics, Cat. No. 90001 and CR-002-01).

In the experiment with Nb.BssSI, molecule N50 was 0.1298 Mbp for molecules above 20 kbp and 0.2336 Mbp for molecules above 150 kbp - with an average label density of 11.8/100 kbp for molecules above 150 kbp. Map rate was 38.9% for molecules above 150 kbp. Effective coverage was 28.2x. In the experiment with DLE-1, molecule N50 was 0.2475 Mbp for molecules above 20 kbp and 0.3641 Mbp for molecules above 150 kbp - with an average label density of 15.7/100 kbp for molecules above 150 kbp. Map rate was 56.4% for molecules above 150 kbp. Effective coverage was 30.6x. Using both Nb.BssSI and DLE-1, label metrics were in line with the manufacturer’s expectations.

### Assembly of optical maps

The *de novo* assembly of the optical maps was performed using the Bionano Access v1.2.1 and Bionano Solve v3.2.1 software. The assembly type performed was the “non-haplotype” with “no extend split” and “no cut segdups”. Default parameters were adjusted to accommodate the genomic properties of the barn swallow genome. Specifically, given the size of the genome, the minimal length for the molecules to be used in the assembly was reduced to 100 kbp, the “Initial P-value” cut off threshold was adjusted to 1 × 10^−10^ and the P-value cut off threshold for extension and refinement was set to 1 × 10^−11^ according to manufacturer’s guidelines (default values are 150 kbp, 1 × 10^−11^ and 1 × 10^−12^ respectively).

A total of 233,450 (of 530,527) NLRS-labelled molecules (N50 = 0.2012 Mbp) were aligned to produce 2,384 map fragments with an N50 of 0.66 Mbp for a total length of 1338.6 Mbp (coverage = 32x). 108,307 (of 229,267) DLE-1 labelled input DNA molecules with a N50 of 0.3228 Mbp (theoretical coverage of the reference 48x) produced 555 maps with a N50 length of 12.1 Mbp for a total length 1299.3 Mbp (coverage = 23x).

### Hybrid scaffolding

Single and dual enzyme Hybrid Scaffolding (HS) was performed using Bionano Access v1.2.1 and Bionano Solve v3.2.1. For the dual enzyme and DLE-1 scaffolding, default settings were used to perform the HS. For Nb.BssSI the “aggressive” settings were used without modification. The NLRS HS had an N50 of 8.3 Mbp (scaffold only N50 = 10.8 Mbp) for a total length of 1,338.6 Mbp (total length of scaffolded contigs = 1,175.3 Mbp) and consisted of 409 scaffolds and 2,899 un-scaffolded contigs. The DLS HS had scaffold N50 of 17.3 Mbp (scaffold only N50 = 25.9 Mbp) for a total length of 1,340.2 Mbp (total length of scaffolded contigs = 1,148.4 Mbp) and consisted of 211 scaffolds and 3,106 un-scaffolded contigs. Dual enzyme HS (incorporating both NLRS and DLS maps) resulted in an assembly with N50 of 23.8 Mbp (scaffold only N50 = 28.4 Mbp) for a total length of 1,351.8 Mbp (total length of scaffolded contigs = 1,208.8 Mbp) and consisted of 273 scaffolds and 2,810 un-scaffolded contigs. During the automatic conflict resolution in the dual enzyme HS, 185 SMRT contigs were cut, as Bionano maps confidently indicated mis-assemblies of the SMRT reads. Conversely, 117 Bionano maps were cut, indicating that the chimeric score did not provide sufficient confidence to cut the assembly based on SMRT contigs. Of 3,872 SMRT contigs, 1,243 (32%) were anchored in the Bionano maps (of which 990 were anchored in both NLRS and DLS maps). 226 and 56 were anchored in NLRS and DLS maps respectively. 2,810 maps could not be anchored at all.

### Purge of haplotigs and final assembly

Notably, all hybrid assemblies were somewhat larger than the expected genome size, and in all cases, the N50 of un-scaffolded contigs was extremely low (0.06 Mbp for the dual enzyme hybrid assembly). We hypothesized that a significant proportion of these small contigs might represent divergent homologous haplotigs that were assembled independently [15]. Similarity searches were consistent with this possibility as almost 95% of the contigs that were not scaffolded in the dual enzyme hybrid assembly showed > 98% identity to scaffolded contigs over 75% of their length or more. These contigs were discarded, resulting in a final assembly (Table 1 and Supplementary Table 1 for detailed statistics) of 1.21 Gbp (N50 = 25.9 Mbp) made up of 273 dual enzyme hybrid scaffolds (N50 = 28.42 Mbp) and 91 un-scaffolded contigs (N50 = 0.0644 Mbp). The final assembly is slightly smaller than the previously estimated genome size (1.28 Gbp) [16]. This potentially reflects an imprecise older estimate, and/or the possibility that some repeated sequences (e.g. centromeric and telomeric low complexity regions) were either collapsed in the initial assembly steps or discarded in the final haplotig purging step described above. The average SMRT read coverage for the genome assembly was 34.15X (implying a theoretical QV of over 40). Supplementary Figure 5 provides a summary of observed sequence coverage depth.

### Assembly metrics for contigs and final scaffolds in our European barn swallow genome

**Table 1:** ^1^ SMRT reads assembled using CANU v1.7 [14]. ^2^ SMRT contigs assembled with CANU and scaffolded using Bionano dual enzyme HS, with haplotigs removed as detailed in the text. *Based on a barn swallow genome size estimate of 1.28 Gbp [16].

|  | SMRT contigs<sup>1</sup> | Final assembly<sup>2</sup> |
| --- | --- | --- |
| Species | *H. r. rustica* |  |
| Starting raw data (Gbp) | 66.4 | 59.6 |
| N50 (bp) | 5189284 | 25954216 |
| N90 (bp) | 85340 | 2002624 |
| Total size (Gbp) | 1.31 | 1.21 |
| Theoretical genome coverage* | 52x | 47x |
| % genome coverage* | 102.6 | 94.5 |
| # of contigs/scaffolds | 3872 | 364 |
| Avg contig/scaffold length (bp) | 338782 | 3334461 |
| Longest contig/scaffold (bp) | 33230000 | 98053015 |

### Annotation of genes and repeats

With respect to mammals, avian genomes generally contain relatively low proportions of repetitive sequences and show strong mutual synteny [17]. This appears to be the case for the barn swallow genome. In particular, 7.11% of the final assembly was annotated as repetitive using WindowMasker [18] and RepeatMasker [19]. The major contributors to annotated repeats were L2/CR1/Rex LINE elements (3.37%), retroviral LTRs (1.59%) and simple repeats (1.56%).

Repeats were soft-masked prior to *de novo* gene prediction using Augustus [20] with *Gallus gallus* gene models. In all, 35,644 protein coding genes were predicted, of which 9,189 were overlapped by more than 30% of their size with repetitive genomic elements. Of the remaining 26,455 predicted protein coding genes, 24,331 harboured a PFAM protein domain (as identified by PfamScan v1.6 [21]). Simple similarity searches based on blastp [22] (with default parameters) suggested that 17,895 of the predicted protein coding genes have a best reciprocal blast hit with gene models derived from *G. gallus* GRCg6a assembly (as available from [23]), while 2,927 of the proteins predicted by Augustus did not show any significant match (e-value <= 1 × 10^−15^, identity > 35%).

### BUSCO genes and phylogenetic reconstruction

Of a total of 4,915 conserved bird Benchmarking with Universal Single-Copy Orthologs (BUSCO) groups [24] sought, 4,598 (93.6%) were complete (and mostly single-copy, 4,521 overall - 92.0%, while 77 were duplicated - 1,6%), 192 (3.9%) were fragmented and 125 (2.5%) were missing. The percentage of contiguously assembled BUSCO genes is consistent with recent results with Anna’s hummingbird (*Calypte anna*) and the Zebra Finch (*Taeniopygia guttata*) [15].

Protein sequences inferred from coding sequences identified by BUSCO v3 as barn swallow orthologs of universal avian single copy genes were aligned to passerine orthologs present in orthoDB v9.0 [25] when all represented passerines had an annotated ortholog. A total of 3,927 protein alignments were generated using the software muscle v3.8.31 [26] with default settings. Software GBlocks v0.91b [27] with default settings apart from allowing gaps in final blocks was used to exclude low quality alignment regions. Trimmed protein alignments were concatenated to produce a supergene alignment with 1,707,664 amino acid positions. Maximum-Likelihood (ML) phylogenetic inference and estimation of aLRT branch support indexes were performed using the software PhyML v3.0 [28], with the LG substitution matrix [29] incorporating 4 variable and 1 invariable gamma distributed substitution rate categories. Distance bootstrap proportions (100 replicates) were estimated using the BioNJ method with the Kimura protein distance correction as implemented in the software SeaView v4.6.5 [26]. ML phylogenetic analysis of concatenated protein sequence alignments yielded a robustly supported topology (Supplementary Figure 6) that is consistent with previous phylogenomic studies [30,31] as well as with gene-level phylogenies [32,33].

### Synteny with the Chicken genome

Alignment of the final assembly with the most recent assembly of the chicken genome (GRCg6a) using D-Genies [34] indicates high levels of collinearity between these two genomes with a limited number of intra-chromosomal rearrangements (Figure 2). The high level of collinearity between independently assembled and scaffolded sequences provides circumstantial support for the quality of both the contigs and the hybrid scaffolds, and is consistent with previous observations of high levels of synteny and minimal inter-chromosomal rearrangements among birds [17].

**Figure 2:**
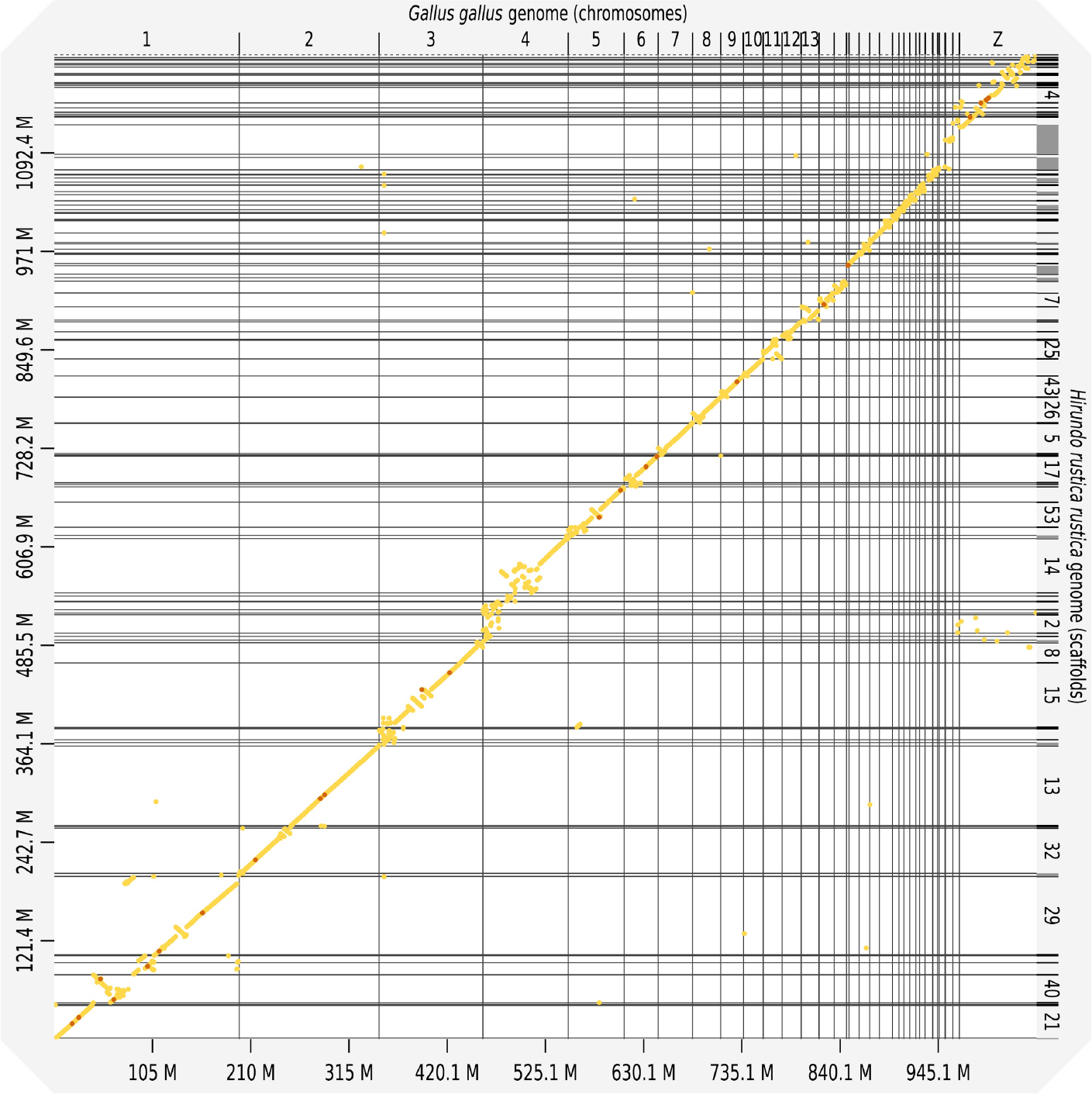
Alignment of the final assembly with the published chromosome-level assembly of the chicken (*G. gallus*) genome GRCg6a using D-Genies [34]. Light to dark yellow dots indicate progressively higher similarity between sequences.

Overall, 90.44% of the chicken assembly can be uniquely aligned to regions in the barn swallow assembly. Table 2 shows for each chicken chromosome (assembly GRCg6a) the number of barn swallow scaffolds aligning uniquely (by best reciprocal BLAST analysis) as well as the percentage of the chicken chromosome involved in alignments. Together with the synteny plot shown in Figure 2, these data indicate that a high proportion (>85%) of most barn-swallow autosomes are likely assembled in less than 10 scaffolds. Indeed several chromosomes are likely assembled as single scaffolds. However, some alignments of chicken chromosomes to the barn swallow assembly are either notably more fragmented or partial. In particular, a large proportion of chicken chromosomes 1 and 4 are represented in unique alignments with the barn swallow assembly. However, for both these chromosomes a number of rearrangements are implied (Figure 2), in line with previous comparisons between the chicken genome and those of other Passeriformes [35–37]. Unambiguous matches between chicken chromosome 16 (2.84 Mb in the chicken assembly GRCg6a, 16 Mb according to flow karyotyping [38]) and the barn swallow assembly were scarce, consistent both with previous reports of difficulties in assembling this chromosome [35,39], likely due to the unusual gene distribution, presence of rRNA repeats, and the polymorphic and often polygenic MHC loci [40] on this chromosome. Similarly, chromosome 31, for which RepeatMasker identified 3.57 Mb (58% of the GRCg6a chromosome assembly) as repeats, was also assembled in a rather fragmented manner in the barn swallow.

Of the sex chromosomes, chicken chromosome Z sequences are well represented, if somewhat fragmented in the barn swallow assembly. The discontinuous assembly of this chromosome is likely related to the widespread presence of repeats [41,42]. For the chicken W chromosome (6.81 Mb in the GRCg6a chromosome assembly, 43 Mb according to flow karyotyping [38]), apparent orthologs of 45 (of 53 single copy genes annotated on the W chromosome of in the *G. gallus* assembly) were identified in the barn swallow genome, although only 46% of the assembled chicken chromosome found best reciprocal BLAST matches. Indeed, Avian W chromosomes are gene-poor and contain long, lineage specific repeats [43,44], complicating both assembly and comparative analyses.

### Alignment between the *G. gallus* GRCg6a and barn swallow genome assemblies

**Table 2:** For each chicken chromosome the number of scaffolds aligning uniquely as well as the percentage of the chicken chromosome involved in alignments are reported.

| Chromosome | N. of uniquely aligned scaffolds | Size in GRCg6a assembly (Mbp) | Covered scaffolds from our assembly |
| --- | --- | --- | --- |
| 1 | 9 | 197.61 | 92.83 |
| 2 | 6 | 149.68 | 88.10 |
| 3 | 7 | 110.84 | 91.70 |
| 4 | 15 | 91.32 | 94.48 |
| 5 | 3 | 59.81 | 89.91 |
| 6 | 2 | 36.37 | 91.87 |
| 7 | 1 | 36.74 | 90.18 |
| 8 | 1 | 30.22 | 90.26 |
| 9 | 1 | 24.15 | 92.65 |
| 10 | 2 | 21.12 | 87.05 |
| 11 | 2 | 20.2 | 89.74 |
| 12 | 3 | 20.39 | 92.40 |
| 13 | 2 | 19.17 | 90.54 |
| 14 | 1 | 16.22 | 91.01 |
| 15 | 1 | 13.06 | 91.62 |
| 16 | 3 | 2.84 | 46.10 |
| 17 | 1 | 10.76 | 93.57 |
| 18 | 3 | 11.37 | 96.86 |
| 19 | 1 | 10.32 | 88.22 |
| 20 | 2 | 13.9 | 92.45 |
| 21 | 1 | 6.84 | 95.03 |
| 22 | 3 | 5.46 | 85.47 |
| 23 | 2 | 6.15 | 87.65 |
| 24 | 1 | 6.49 | 92.78 |
| 25 | 1 | 3.98 | 90.50 |
| 26 | 1 | 6.06 | 94.06 |
| 27 | 3 | 8.08 | 96.75 |
| 28 | 3 | 5.12 | 94.61 |
| 30 | 13 | 1.82 | 72.81 |
| 31 | 4 | 6.15 | 28.14 |
| 32 | 6 | 0.73 | 95.35 |
| 33 | 5 | 7.82 | 92.63 |
| W | 5 | 6.81 | 45.90 |
| Z | 34 | 82.53 | 89.82 |

## Conclusion

Short-read NGS (now known as Second Generation Sequencing, or SGS) technologies have allowed the production of cost-effective genome drafts for many birds and other vertebrate species [30,45,46]. However, the reduction in genome sequencing costs has typically come at the price of compromises in contiguity and accuracy of assemblies with respect to earlier efforts based on Sanger reads and extensive physical mapping [47]. Many limitations of SGS-based assemblies stem from the occurrence of long sequence repeats. In many animal species, transposons are frequently located in introns [48] and the presence of large gene families of closely related paralogs can lead to the existence of long “genic” repeats. Accordingly, even apparently contiguous genic regions can feature juxtaposition of paralogous gene fragments [15]. Given the inception of large scale sequencing initiatives aiming to produce genome assemblies for a wide range of organisms [49–52], it is critical to identify combinations of sequencing and scaffolding approaches that allow the cost effective generation of genuinely high-quality genome assemblies [10]. While exhibiting higher rates of single-base errors than some SGS approaches, TGS technologies, including SMRT sequencing, offer read-lengths unparalleled by SGS or Sanger sequencing [53]. Moreover, recent and ongoing improvements in TGS methods are rapidly reducing the “per-base” cost of TGS data compared to that of SGS. On the other hand, as an alternative to scaffolding with long insert mate-pairs [54] or to chromatin proximity ligation sequencing [55], contiguity and accuracy of long-read-based assemblies can be further improved by optical mapping. This relies on nanoscale channels that can accommodate thousands of single, ultralong (>200 kbp) double-stranded DNA filaments in parallel, subsequently stained to recognize specific 6-7 bp long motifs [56]. The combination of long reads and optical maps has already proven invaluable to produce high-quality genome assemblies, even in the case of particularly complex genomes [57]. Here, using only SMRT sequencing and Bionano optical maps we have produced a high-quality and contiguous genome for the barn swallow. With respect to a previously reported SGS-based assembly of the American barn swallow genome using a comparable amount of raw data [2], even the contigs generated from long-read sequencing alone show a 134-fold increase in N50, similar to the increase recently obtained for the Anna’s hummingbird genome using the same technologies [15]. Furthermore, the 1.6 fold change in scaffold N50 attained by Bionano NLRS hybrid scaffolding before removal of haplotigs is comparable with results obtained by other genome assemblies that have employed this method [58]. Strikingly, the new DLS method greatly outperformed the NLRS system, providing a 3.3 fold increase of N50 (before removal of haplotigs).

Moreover, incorporation of both labelling systems into the hybrid scaffolding yielded a final assembly showing 5-fold improvement of the N50 with respect to the original SMRT assembly, simultaneously providing “independent” validation of many scaffold junctions. We note that the presence of numerous microchromosomes in avian genomes restricts the final N50 value potentially attainable for the assembly, as for example the fully assembled karyotype of the chicken genome assembly (GRCg6a) would have an N50 of ~ 90 Mbp. Yet, after removal of putative haplotigs, our genome assembly contiguity metrics meet the high standards of the VGP consortium “Platinum Genome” criteria (contig N50 in excess of 1 Mbp and scaffold N50 above 10 Mbp) [10]. Accordingly, we believe that the data presented here, while attesting to the effectiveness of SMRT sequencing combined with DLS optical mapping for the assembly of vertebrate genomes, will provide an invaluable asset for population genetics and genomics in the barn swallow and for comparative genomics in birds.

### Re-use Potential

Future directions for the barn swallow genome will include further scaffolding using a G10K VGP approach, the phasing of the assembly to generate extended haplotypes, a more thorough gene annotation using RNA/IsoSeq sequencing data, detailed comparisons with the genome of the North American subspecies, *H. r. erythrogaster*, studies on the genomic architecture of traits under natural and sexual selection, and the re-evaluation of data from population genetics studies conducted in this species (as it was shown that the availability of a high quality genome may change the interpretation of some results), as well as characterization of the epigenetic landscape.

## Availability of supporting data

The data sets supporting the results of this article will available in the GenBank repository upon acceptance, under Bioproject PRJNA481100.

## Authors’ contributions

G.F, N.S., A.B.-A., L.G., D.S.H, M.C. and L.C. conceived the project and designed the experiments; G.F. performed DNA extraction and quality control; M.C. carried out CANU assembly, gene and repeat annotation. D.S.H., M.C. and L.G. performed other bioinformatics analyses; L.P. conducted the optical mapping; K.J.F. produced the hybrid scaffolds; G.F., D.S.H, M.C., N.S. and L.C. drafted the manuscript. All authors edited and contributed to the manuscript.

## Acknowledgements

We thank Manuela Caprioli for support in field work, sample collection, DNA extraction and quality control as well as Dr. Elena Galati for support in PFGE quality control. We also thank The Functional Genomics Center of Zurich, where SMRT sequencing and optical mapping were carried out, and particularly Andrea Patrignani for SMRT sequencing. We are thankful to the Genome 10K Council and to all members of the Consortium for the support in obtaining early access to the DLS technology. We are particularly thankful to the G10K Chair Prof. Erich Jarvis, also for his helpful comments to the manuscript. We thank Chiara Scandolara for the barn swallow picture used for Figure 1. We acknowledge the support of ELIXIR-IT and CINECA (HPC@CINECA) for provision of computational resources for SMRT read assembly. We thank the reviewers who helped us to considerably improve the first version of this manuscript.

## Ethics approval

The blood sample used to generate the genomic data derived from a minimally invasive sampling on a single individual. Appropriate consent was obtained from the local authorities (Regione Lombardia).

## Additional files

**Supplementary Figure 1(Supplementary Figure 1.png).**
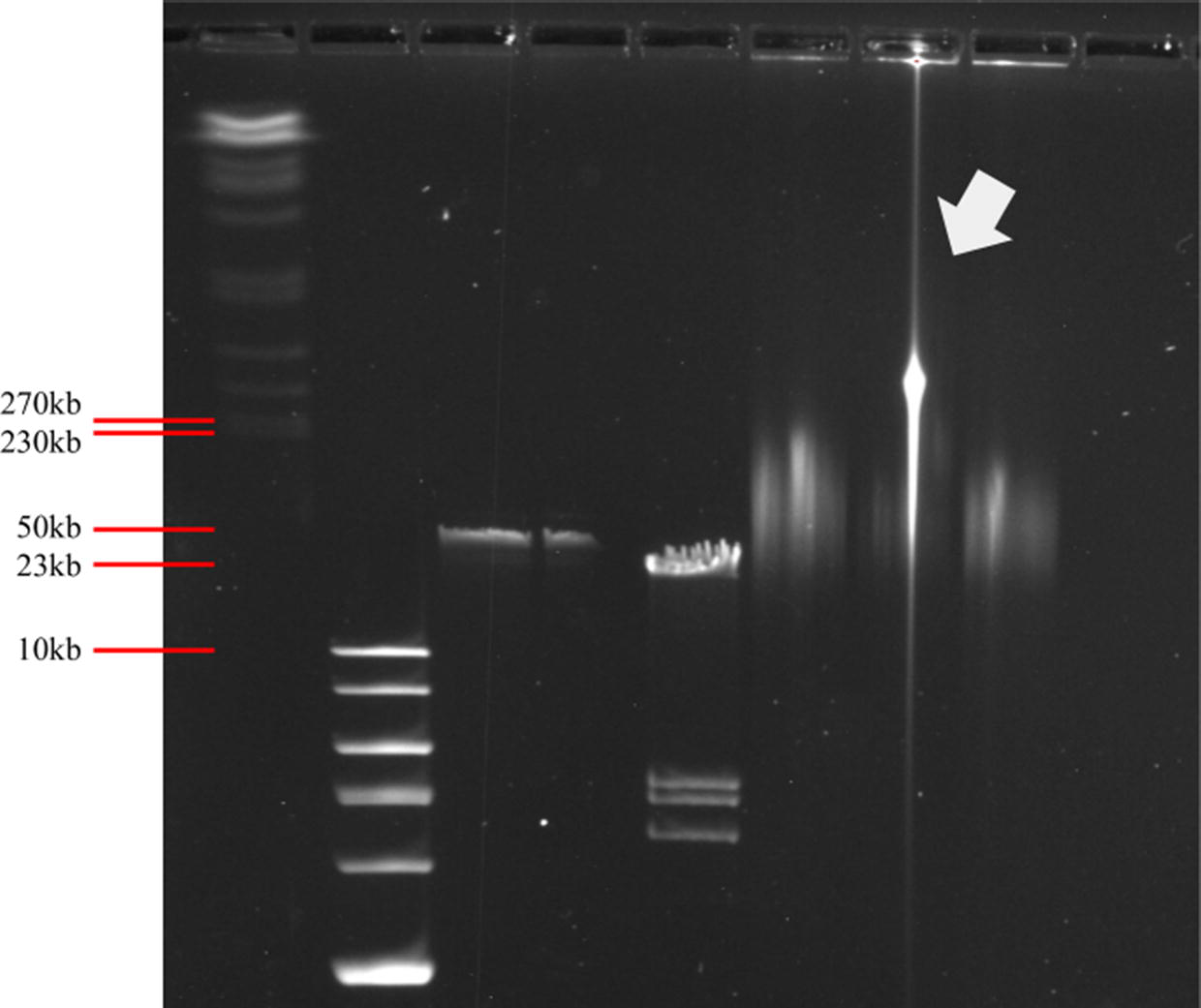
PFGE on a 1x agarose gel run for 18 hours at 160 mV. The two lowest overlapping bands in lane 1 represent yeast chromosomes of 230 kbp and 270 kbp, respectively. Lane 2 contains 1kb DNA ladder (highest 10 kbp), lane 3 and 4 the undigested lambda phage (50 kbp) and lane 5 digested lambda (upper band 23 kbp). Lane 7 contains the sample used in the study.

**Supplementary Figure 2(Supplementary Figure 2.tif).**
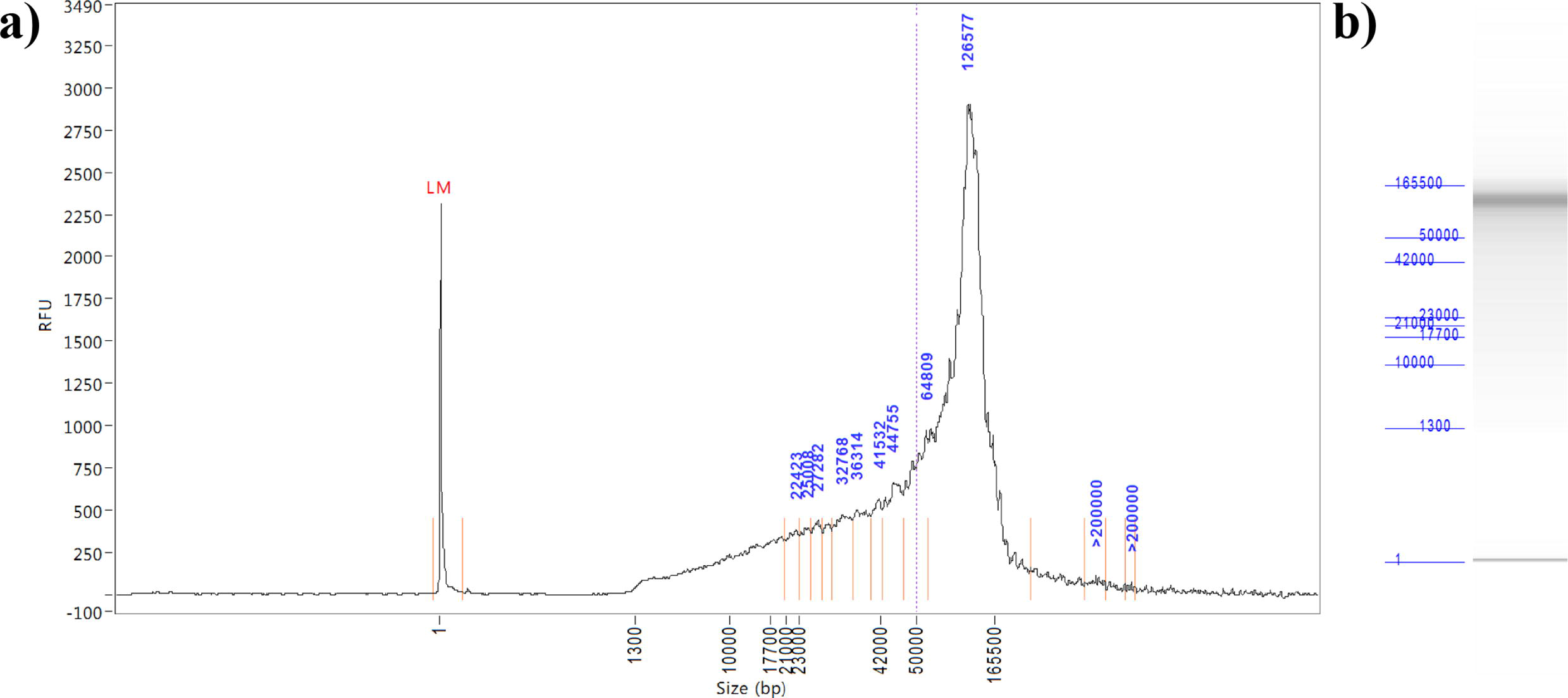
FEMTO Pulse capillary electrophoresis results from software PROSize Data Analysis (AATI) for the DNA sample used in the study. a) Quantity by fragment size plot. The software algorithm identifies the peaks of major fluorescence change (defined within the range of 2 orange bars) and assign a size value to them (blue numbers). The purple dashed line represents the 50 kpb cutoff. RFU = Relative Fluorescence Unit. LM = Lower Marker. b) Virtual gel. Note that DNA > 200 kbp is above the detection range of the instrument and is conventionally labelled as > 200 kbp.

**Supplementary Figure 3(Supplementary Figure 3.png).**
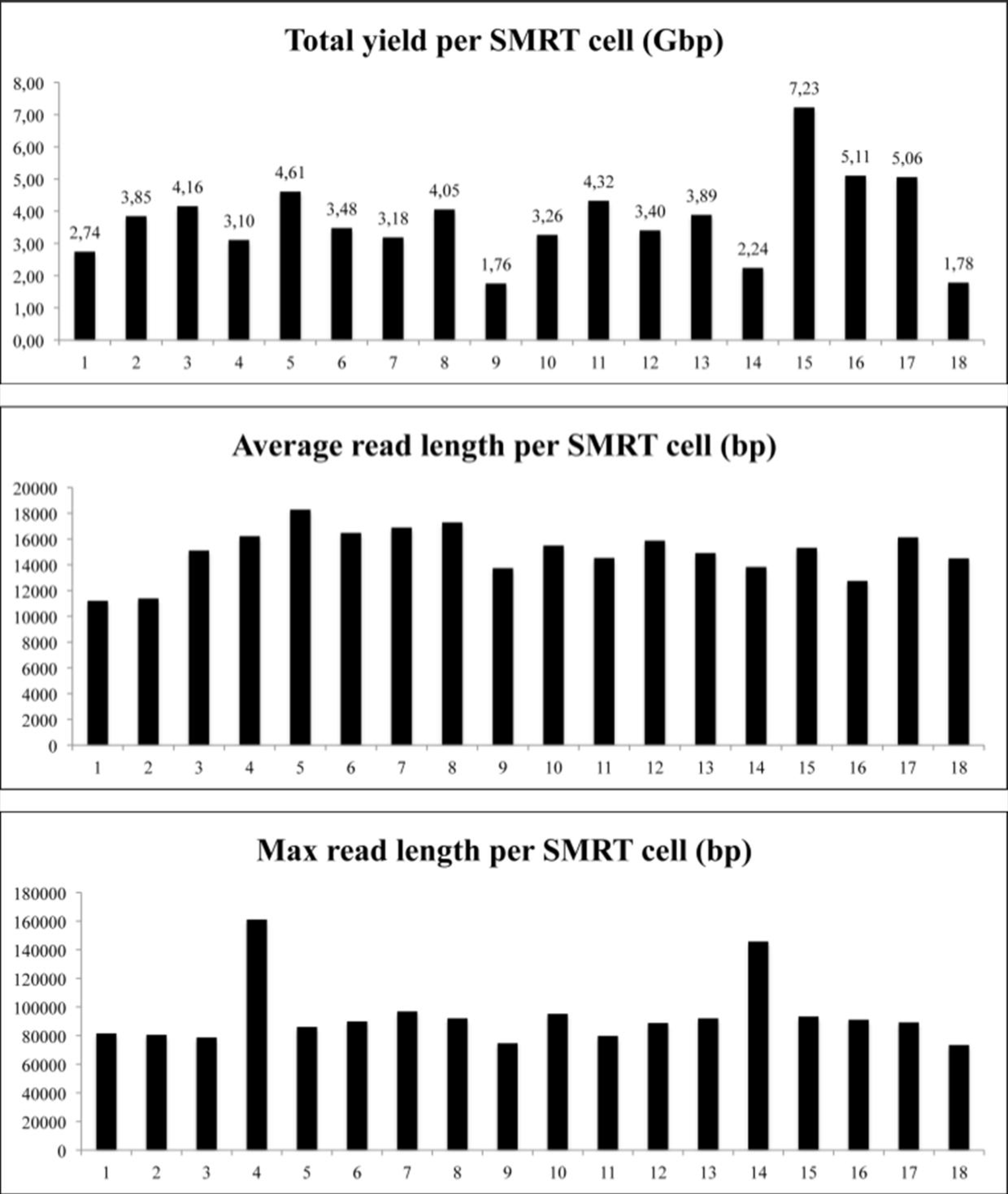
Summary statistics for each SMRT cell employed.

**Supplementary Figure 4(Supplementary Figure 4.png).**
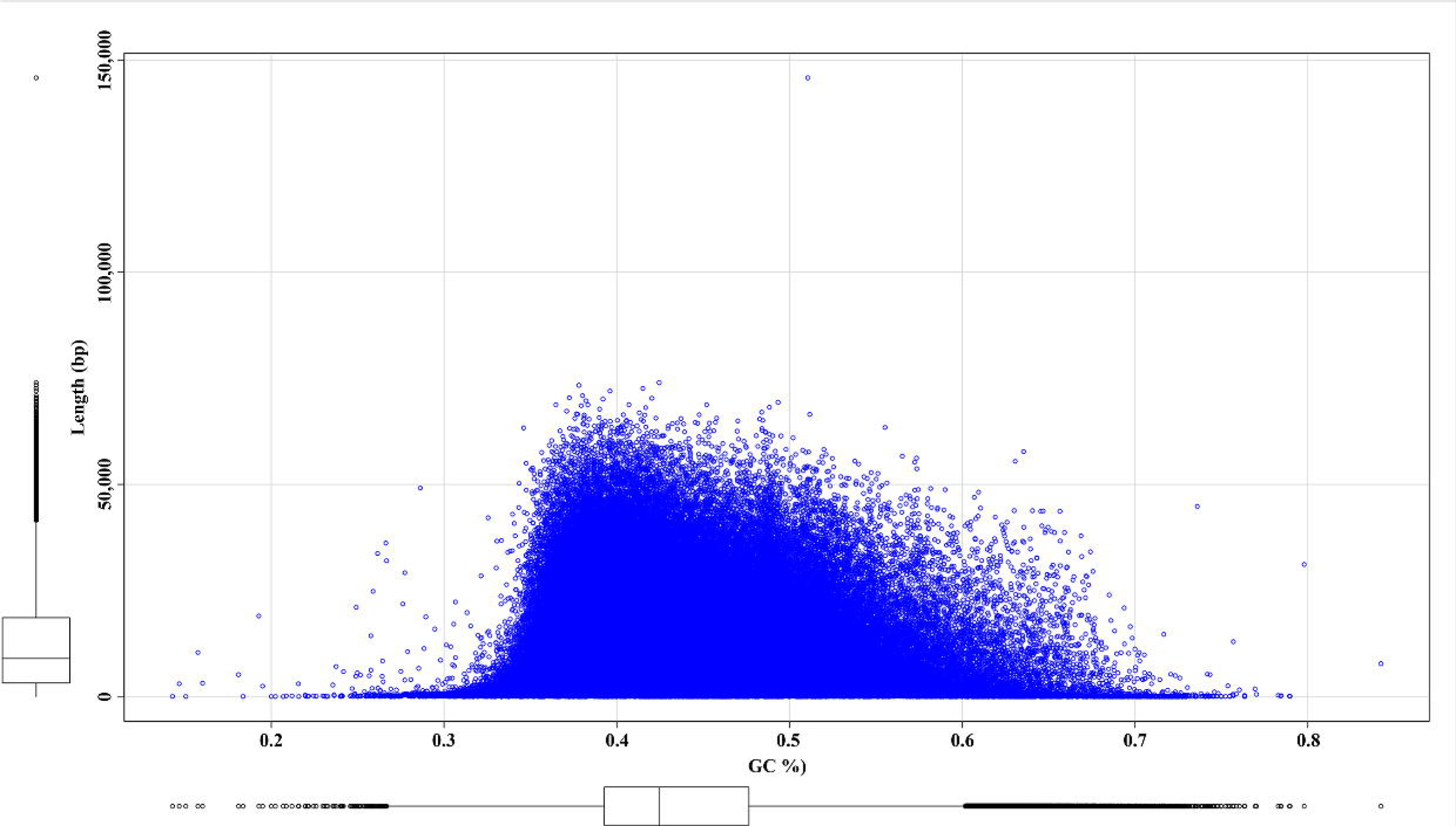
GC content distribution in all sequence reads after CANU trimming.

**Supplementary Figure 5(Supplementary Figure 5.tif).**
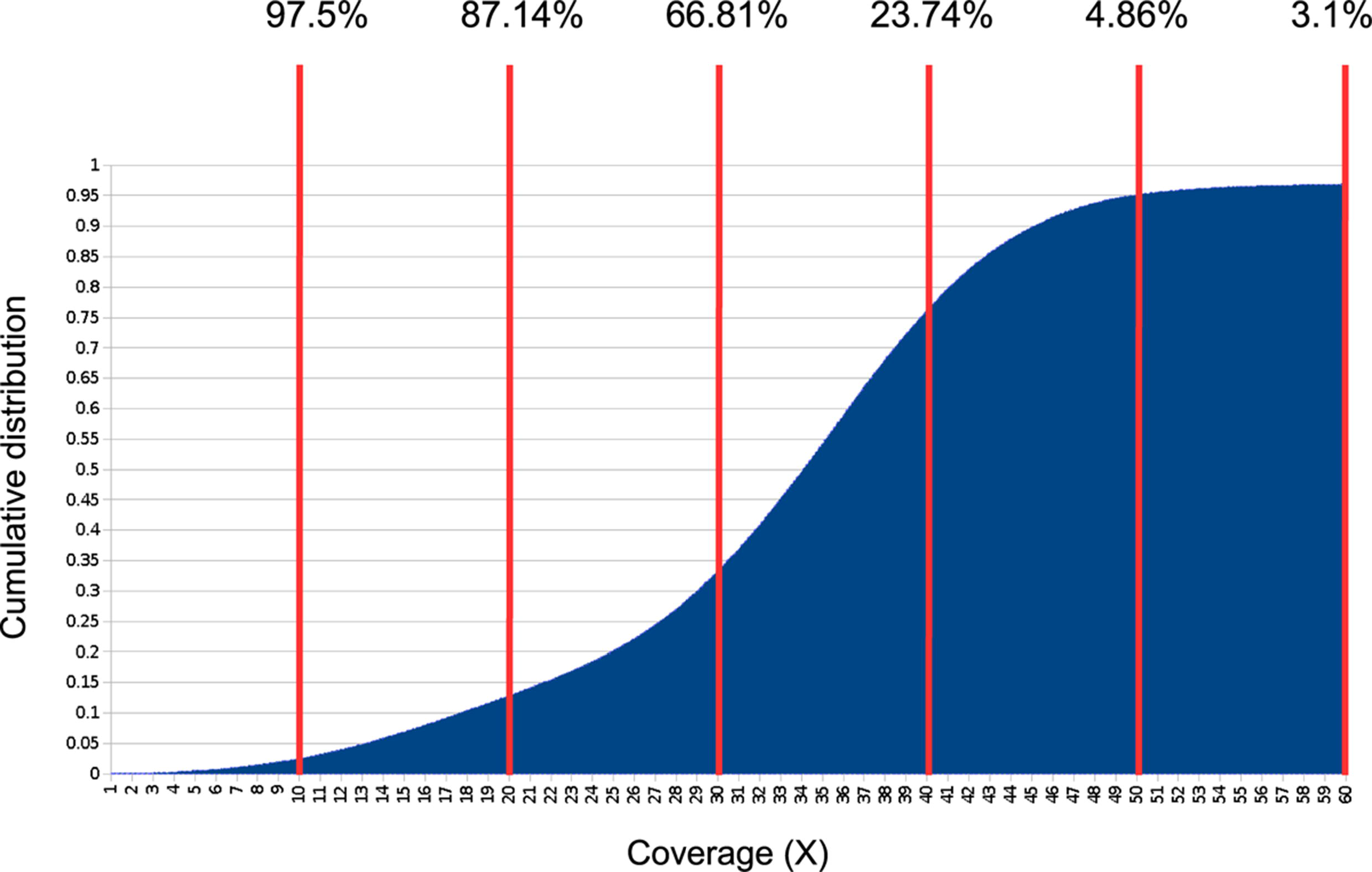
Cumulative coverage distribution of the final (de-haplotyped) assembly of the barn swallow genome. Coverage is indicated on the X axis. Red lines are used to display the proportion of the genome covered by more than 10, 20, 30, 40, 50 or 60 reads respectively.

**Supplementary Figure 6(Supplementary Figure 6.eps).**
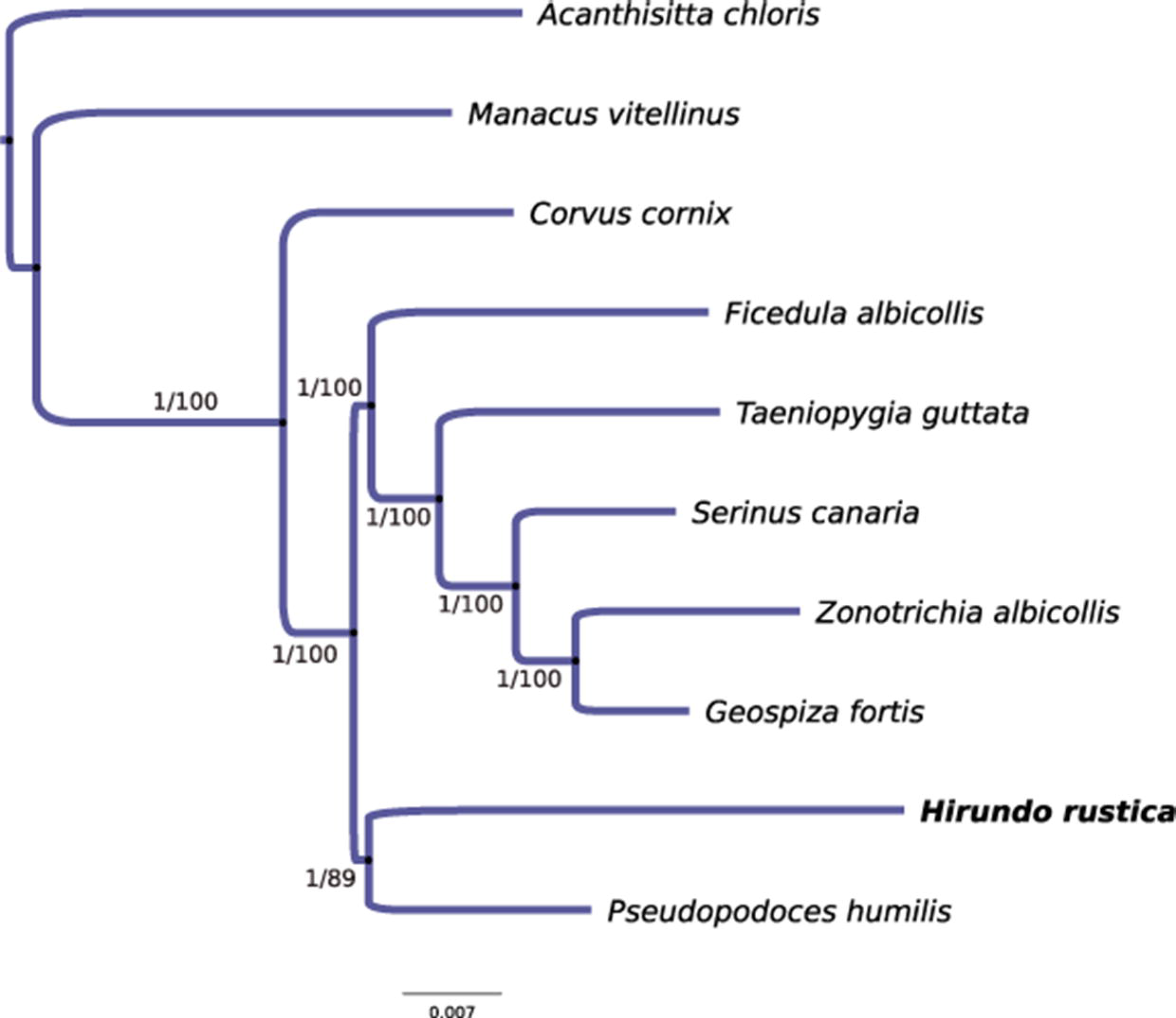
Maximum likelihood phylogenetic tree based on a multiple alignment of 3,927 gene orthologs in passerine species. The scale bar indicates inferred changes per site, aLRT support values and neighbor joining bootstrap values (100 replicates) are shown on branches.

<u>Supplementary Table 1</u> (Supplementary Table 1.xlsx)

Comparison of assembly metrics for contigs and scaffolds between different assemblies. In hybrid scaffolds, the first column refers to assemblies including the un-scaffolded contigs while the second column only includes scaffolded contigs metrics. The estimated genome size of 1.28 Gbp is from [16]. Average gene size was estimated according to the latest available annotation of the *G. gallus* genome (GRCg6a).

## List of abbreviations

DLS: Direct Label and Stain
HMW: High Molecular Weight
HS: Hybrid Scaffold
NGS: Next Generation Sequencing
NLRS: Nick, Label, Repair and Stain
N50: the shortest sequence length at 50% of the genome
N90: the shortest sequence length at 90% of the genome
PFGE: Pulsed Field Gel Electrophoresis
QV: Quality Value
SGS: Second Generation Sequencing
SMRT: Single Molecule

